# Simultaneous detection of RNA modifications using *In Vitro* Transcription and Direct RNA Nanopore sequencing

**DOI:** 10.1101/2024.10.22.619587

**Authors:** Logan Mulroney, Lucia Coscujuela Tarrero, Paola Maragno, Carmela Rubolino, Simone Maestri, Mattia Furlan, Matteo Jacopo Marzi, Tom Fitzgerald, Tommaso Leonardi, Mattia Pelizzola, Ewan Birney, Francesco Nicassio

## Abstract

RNA modifications are critical for transcript function and regulation, yet detecting these modifications transcriptome-wide and at isoform-level resolution remains technically challenging. Here, we present a robust *in vitro* transcription (IVT)-based strategy coupled with direct RNA nanopore sequencing (dRNA-seq) to detect a wide spectrum of endogenous RNA modifications without requiring prior knowledge of modification types or modification-specific biochemical assays. We generated an IVT modification-free reference transcriptome from K562 cells and used Nanocompore to compare it to the native RNA. We detected 26,619 modification sites across 2,520 isoforms from 1,766 genes. We used motif and annotation-based inference to identify at least eight distinct RNA modifications, with m6A and m5C being the most prevalent. Importantly, we uncovered non-random co-occurrence of m6A and m5C on both the same transcripts and a subset of the same molecules, suggesting potential combinatorial regulation. Furthermore, RNA modification patterns were often isoform-specific, pointing to a link between the epitranscriptome and alternative splicing. This approach also revealed previously underexplored modification patterns in mitochondrial mRNAs, suggesting broader regulatory complexity than previously appreciated. Our study provides a survey of RNA modifications across the transcriptome, demonstrating the utility of *in vitro* transcription coupled with direct RNA nanopore sequencing to simultaneously detect multiple modifications without the need for additional independent biochemical assays, enabling future investigations into the dynamics, coordination, and functional consequences of RNA modifications across different biological contexts.

## Background

The study of gene expression and regulation is essential to understand the mechanisms underlying cellular processes and diseases (1). Transcriptome analysis provides a comprehensive view of gene expression by detecting and quantifying RNA transcripts; depending on the details of the RNA preparation, one can also determine alternative splicing patterns. Cell metabolism is influenced at many levels (2), including transcription regulation (3), splicing (4), and translation (5). The biological function of RNA molecules is determined by their sequence, structure, and chemical modifications (6,7). RNA modifications are chemical alterations of nucleotides that often change the properties of RNA molecules, including their stability, structure, cellular localization and ultimately function (7–9). Despite the extensive and now routine use of RNA-seq experiments to assess RNA levels and alternative splicing, it has been challenging to assess RNA modifications in individual transcripts at single-nucleotide resolution. Although there is considerable evidence for the presence of RNA modifications and some understanding of their generic role, rarely are the RNA modifications resolved to individual transcripts or the role of individual RNA modifications on specific transcripts.

RNA modifications can be broadly classified into two categories: nucleoside modifications and base modifications. Nucleoside modifications involve the addition or removal of functional groups to the ribose or deoxyribose sugars of nucleotides, while base modifications involve the modification of the nitrogenous bases of nucleotides (10). Among base modifications, the most abundant mammalian internal mRNA modification is N6-methyladenosine (m6A), which has been shown to regulate mRNA stability, splicing, and translation (11–13). Another commonly identified RNA modification is 5-methylcytosine (m5C), which is found in tRNA, rRNA, and mRNA, where it is involved in translation and RNA folding (14,15). Pseudouridine () is the most abundant mammalian RNA modification that affects RNA structure and stability (16), and 2’-O-methylation (Nm) is found in rRNA, snRNA, and mRNA, where it is involved in ribosome biogenesis and splicing (17,18). Methyl-Pseudouridine incorporation has been critical for the deployment of mRNA vaccines (19), allowing the rapid development of vaccines for infectious disease (20,21) and the potential for individualized cancer vaccines. To date, approximately 170 naturally occurring RNA modifications have been identified (22) and the community has expressed enthusiasm in systematically assessing their presence and relationships to other biological features (23). Multiple chemical modifications often coexist on a single transcript, potentially interacting with binding partners in distinct yet interrelated ways. However, current methodologies typically focus on detecting one specific modification at a time, making it challenging to capture how multiple modifications might jointly affect RNA function. This gap in technology and analytical capacity underscores a significant hurdle in comprehensively understanding post-transcriptional regulation.

RNA modifications can be detected using a variety of methods, often through the results of chemical or enzymatic reactions that are specific for the RNA modification or modifications of interest (24–26). In recent years, direct RNA nanopore sequencing (dRNA-seq) has emerged as a promising approach for detecting multiple RNA modifications (27). dRNA-seq processes RNA without any chemical intermediates, which doesn’t erase the RNA modifications during sequencing (28). Thus, dRNA-seq has the potential to detect all RNA modifications which change the raw ionic current signal, without additional biochemical assays for particular RNA modifications. This detection happens in the context of individual molecules, thus it is possible to detect RNA modification co-occurrence and determine its relationship to the precise transcript structure.

Recently, there have been many studies that used dRNA-seq to investigate RNA modifications, some of which have developed new bioinformatic tools to detect the modified RNA bases (29–31). Some of these tools can detect the RNA modifications from a single sample using characteristic ionic current shifts caused by the modification using training data of known modifications, while others require comparing the sample with a sample that is devoid of the modification(s) of interest (29).

The methods that detect RNA modifications from a single sample typically use neural networks trained to detect particular RNA modifications (29). These tools require laborious sample preparations of specific modifications to generate robust training data for the models. The two-sample comparison methods typically employ statistical tests to find significant differences at each position between the two sample’s distributions of ionic current features or systematic basecalling errors (29). Single-nucleotide precision is challenging for these comparative methods because the ionic current for any state is derived from 5-9 nucleotides that occupy the sensitive region of the nanopore at each step of the motor regulated translocation (32) and the RNA modifications can alter the ionic current for any of these positions. Despite this progress, the need for robust training data or precisely matched pairs of samples can limit the applicability of these methods, particularly for less-characterized modifications.

Trained model methods generally outperform the two-sample comparative approaches for detecting the target RNA modification (33). However, due to the challenges generating the training data, robust trained models are only available for a few well-characterized RNA modifications (29). Comparative methods are thus invaluable when trained models are not available, or multiple RNA modifications are involved. These methods use a reference sample that can be generated using two distinct strategies: altering the deposition or removal of a target RNA modification through genetics or environmental stimulation (34,35); or by generating a synthetic copy of the target RNA without any modified nucleotides (36,37). Although useful for synthesizing short oligonucleotide polymers, phosphoramidite chemistry is not practical for recreating an entire transcriptome (38). Thus, to create a synthetic copy of a sample specific RNA transcriptome, *in vitro* transcription (IVT) is more practical to create a large-scale, modification-free reference of a given biological sample’s transcriptome. Transcriptome-wide IVT synthesis has been used with short-reads as a negative control for antibody based detection of m6A (37). Additionally, this method was recently adapted to dRNA-seq for nanopore-based modification analyses (39,40), underscoring the utility of such an approach for comparative studies of RNA modifications.

Here, we describe a practical protocol for generating IVT-derived RNA at scale, paired with direct RNA nanopore sequencing, to identify putative RNA modifications in a human cell line (K562) by comparing native vs. IVT samples with Nanocompore (35), an analytical framework that finds sites significantly different between two samples by employing a Gaussian Mixture Model to cluster the ionic current in two dimensions for every read, position-by-position (35). We then draw upon existing annotations (RMBase, and writer motifs annotations) to tentatively assign specific modification types (e.g., m6A, m5C) to certain sites. Our analysis of K562 transcripts using this approach identified 26,619 putative modification sites across 2,520 isoforms from 1,742 genes. We were able to classify 8 different RNA modifications. Approximately 56% of these sites were consistent with m6A, and ∼15% were consistent with m5C based on known motifs and database annotations. A fraction of putative m6A and m5C sites were found in proximity (within 25 nt) on the same transcripts, and in a few cases, they appeared to co-occur in the same molecule for specific genes (notably *KRTCAP2*). We also observed that some modifications appeared differentially present in distinct transcript isoforms of the same gene, which could reflect isoform-dependent regulation. Collectively, our results highlight the feasibility of using IVT-based reference samples and dRNA-seq to detect multiple modifications simultaneously. Furthermore, by integrating the IVT approach with long-read sequencing, we successfully captured full-length transcripts and uncovered previously hidden differential isoform modifications, revealing variation in chemical marks that shorter reads often miss, enabling a deeper insight into how these subtle modifications influence RNA regulation.

## Results

### RNA in vitro transcription optimization

The T7 RNA polymerase is commonly used for IVT reactions. To synthesize a sample-specific IVT transcriptome, the T7 promoter must be present on the 5′ end of a double stranded DNA template sequence for each desired transcript. This can be achieved by using a Template Switching Oligonucleotide that contains the T7 promoter sequence during cDNA synthesis with a template switching reverse transcriptase (**Figure 1A**). We tested several conditions for each step of the IVT RNA synthesis procedure and found that the the NEB template switching kit combined with incubating the IVT reaction at 42°C, and in vitro polyadenylating the IVT RNA for 30 minutes were optimal conditions (see **Supplementary Results** and **Supplementary** Figures 1 **& 2** and **Supplementary Tables 1 & 2** for details on our optimization).

**Figure 1:**
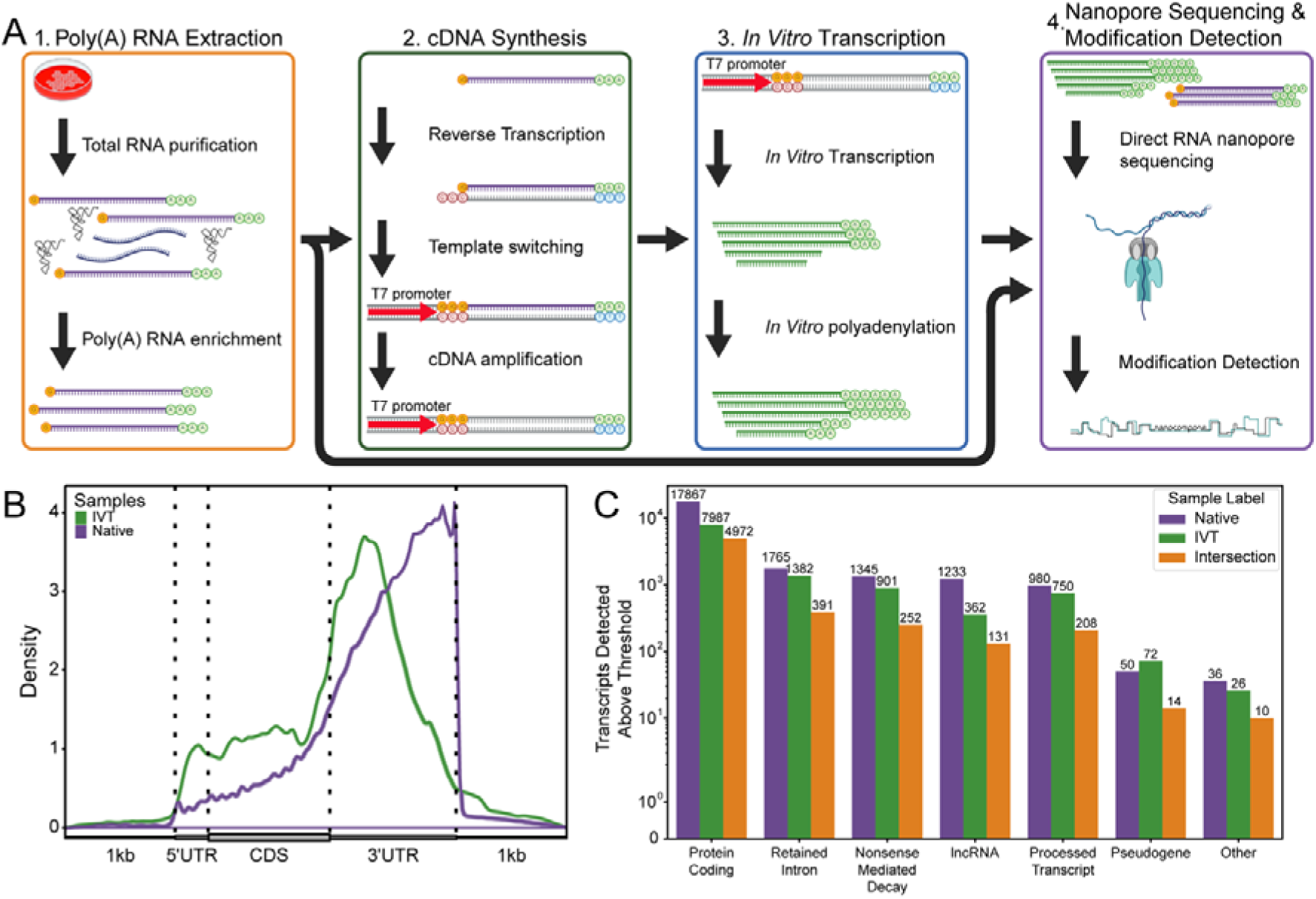
RNA In Vitro Transcription protocol and sequencing coverage: (A) Protocol diagram starting from RNA extraction and resulting in direct RNA nanopore sequencing of the native sample compared with the in vitro transcribed copy. (B) Metagene plot of the direct RNA nanopore sequencing coverage for the Native (purple) and IVT (green) samples across annotated mRNA transcripts. (C) Number of unique transcripts that were detected with more than 30 reads separated by biotype for the Native (purple), IVT (green), and by both the native and IVT samples (orange).

Using our protocol, we obtained 5.4 and 2.9 million pass reads from two IVT samples sequenced on PromethION flowcells. We found that 36.1% of the primary reads were reverse alignments when aligned to the transcriptome reference, which mostly initiated near short poly(C) sequences. These reads are likely internally primed template switching oligos that initiate T7 RNA synthesis from the reverse strand. Accordingly, we removed such sequences and obtained 2.8 and 1.5 million forward primary alignments to the transcriptome reference, respectively (**Supplementary Table 3**). From two Native samples, we obtained 9.1 and 8.7 million pass reads resulting in 7.7 and 7.3 million forward primary alignments to the transcriptome reference, respectively (**Supplementary Table 3**). As expected, due to the 3’-5’ sequencing of RNA molecules requiring a polyA tail, the read coverage for the forward primary alignments were biased towards the 3′ end of most transcripts (**Figure 1B**). However, the IVT coverage was more even across the CDS and 5′ UTR compared to the Native coverage, likely due to the incomplete transcription products getting *in vitro* polyA tailed (**Figure 1B**).

The IVT reads do not need to perfectly match the length profile or detected abundance as the native dRNA-seq reads. Rather, the IVT reads need to cover enough of the transcriptome to provide the representative ionic current signal for each unmodified kmer in the sample. For example, Nanocompore requires 30 reads to cover each position by default (35). The unique transcripts identified by the IVT were primarily a subset of the Native sample, with some differences in per-transcript coverage reducing the total number of possibly analyzed transcripts (**Figure 1C**).

### RNA modification detection

Nanocompore is a comparative tool that was found to have higher specificity than other comparative tools, especially for high-coverage samples (33). Moreover, it allows extracting modification probabilities at base pair level resolution and read-level, a feature that is highly desirable for assessing the co-occurrence of multiple modifications on the same RNA molecule (35).

We used the MinION platform to iterate on the IVT protocol development, and the PromethION to increase throughput and cover more of the transcriptome. There are hardware differences between the MinION and PromethION platforms that can cause differences in how the electric signals are detected and stored. To test if we could combine the data from both platforms for RNA modification detection we used Nanocompore with the PromethION IVT samples and either the MinION or PromethION Native samples. Indeed, we found that the modifications detected were different depending on which platform was used for the Native samples (**Supplementary** Figure 3A **& C**). These differences remained after balancing the coverage between the two platforms (**Supplementary** Figure 3B **& D**). We therefore decided to use only the PromethION data for both IVT and Native samples for the rest of our tests.

From the 4261 transcripts, there were 2.97M total positions analyzed by Nanocompore (see methods and **Supplementary table 4**). We found 26619 significant positions in 2520 transcripts from 1766 genes (**Supplementary table 4**). The number of significant sites per transcript ranged from 1 to 149, with the largest population of transcripts containing only a single modified position (**Figure 2A**).

**Figure 2:**
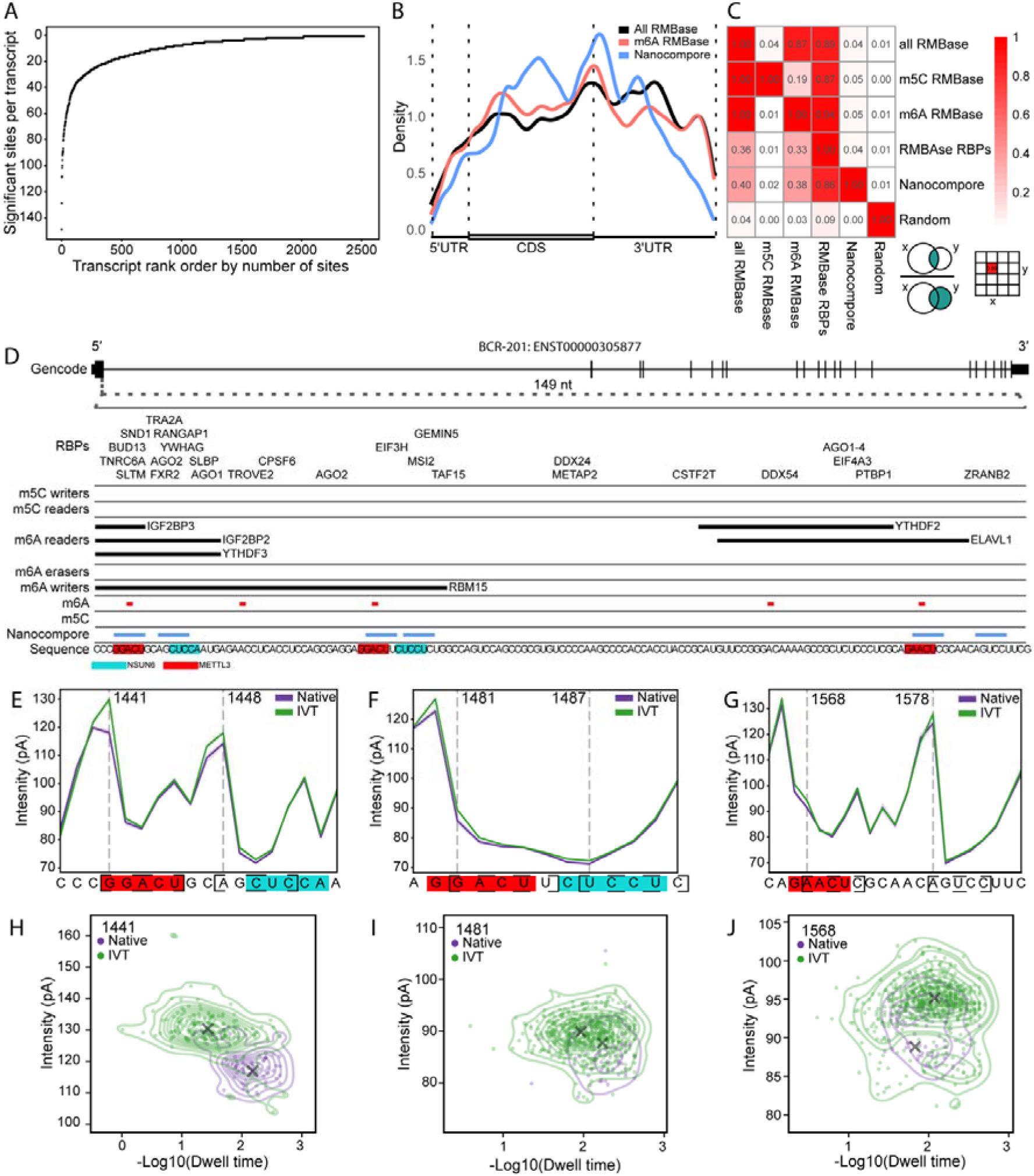
Detected RNA modifications. (A) Scatterplot showing the number of Nanocompore significant sites per transcript isoform, ranked by abundance. Each point represents a transcript; the y-axis indicates the number of significant sites, and the x-axis their rank. (B) Metagene plot of Nanocompore significant sites (blue), RMBase3.0-annotated m6A sites (red), and all RMBase3.0 modifications (black). Vertical dashed lines separate the 5′ UTR, CDS, and 3′ UTR. (C) Heatmap showing overlap between Nanocompore significant sites and RMBase3.0 features. Each cell indicates the fraction of overlapping bins between rows and columns. (D) IGV trace of a 149-nt region from BCR transcript ENST00000305877, showing 6 of 9 Nanocompore significant sites. The top 8 rows display RMBase3.0 annotations, including RNA-binding proteins (RBPs), m A/m C effectors, and modification sites. The Nanocompore row highlights significant sites, and the bottom row shows transcript sequence. Red-highlighted sequence corresponds to METTL3’s DRACH motif; cyan denotes NSUN6’s CUCCA or CUCCU motifs. (E) Line plot of mean ionic current intensity for native (purple) and IVT (green) reads across the first significant site pair. Shaded areas represent one standard deviation. Vertical dashed lines mark significant positions 1441 and 1481; the dashed box shows the k-mer at the nanopore-sensitive region. (F) and (G) are analogous line plots for two additional regions. (H) Two-dimensional density plot of ionic current (y-axis) versus –log₁₀ dwell time (x-axis) at position 1441, comparing native (purple) and IVT (green) reads. GMM cluster centers are indicated by black Xs. (I) and (J) show similar two-dimensional distributions for positions 1481 and 1568, respectively.

The IVT sample is composed entirely of canonical nucleotides, thus all types of RNA modifications can potentially be identified. However, from this analysis alone, the identity of these modifications is unknown. The RNA modification identity can be inferred by matching the sequence context with known motifs for RNA modification writing enzymes or by comparing the sites identified with Nanocompore to RNA modification annotations, such as those compiled by RMBase3.0 (41). As RNA modifications are dynamic, not every RNA modification annotated site exists in every cellular condition, and not every recognition motif is modified by these enzymes (42). Akin to transcript annotations, we can use these RNA annotations to infer the detected modification identity, but not use the annotations as proof of existence in our samples. We identified eight different types of RNA modifications when compared to the annotations, of which 41.13% were m6A, 8.94% were m5C, 5.17% were pseudouridine, 1.17% were m7G, 0.83% were m3C, 0.3% were m1A, 0.27% were Nm, and 0.04% were inosine (**Supplementary Table 5**). We compared the distribution of the Nanocompore significant sites with those annotated by RMBase across a metagene representation (see methods), and found that both the Nanocompore sites and RMBase annotations have a peak near the stop codon (**Figure 2B**). It’s been previously documented that m6A is predominantly detected in the 3′ UTR adjacent to the stop codon (43), as can be seen when only the m6A annotations are plotted in the metagene (**Figure 2B**). The Nanocompore sites had more density in the CDS than what is annotated by RMBase. Considering that m6A is the dominant modification annotated in RMBase and Nanocompore can detect many other modifications which may be distributed differently across the transcripts, this suggests that the IVT-based analysis identified sites of RNA modifications that are different from m6A. Furthermore, we compared the inferred m6A sites to the HEK293 GLORI annotated sites (**Supplementary** Figure 4) and found similar levels of overlap as GLORI compared with miCLIP2 (44).

To more precisely evaluate the overlap between the annotations and the Nanocompore significant sites, we binned the coding regions of the genome into 25 nucleotide wide non-overlapping windows and looked at the bins that were shared between the two sets (see methods, **Figure 2C** & **Supplementary figure 5**). We found that 38% of the bins with Nanocompore significant sites overlapped with a bin containing an RMBase m6A site. When the Nanocompore positive bins were compared to all RNA modifications annotated by RMBase, there were only an additional 2% of the bins that overlapped (40% overlap in total). However, 89% of the Nanocompore positive bins overlapped with an RMBase annotated RNA binding protein associated with modifications. This represents a 9-fold increase in overlaps when compared to randomly shuffling the Nanocompore significant sites (**Figure 2C**). The data and analysis support using IVT baselines for an unbiased calling of the epitranscriptome.

### Coordination between RNA modifications

To understand the possible RNA modifications identified by Nanocompore, we examined the sites found in BCR, a gene typically found mutated in leukemia, the source tissue for K562 cells (45). We found nine Nanocompore significant sites in BCR, six of which were found in the first exon (**Figure 2D**). Surprisingly, these six significant sites appeared in pairs, where the first site of each pair overlapped with both m6A sites annotated by RMBase and the DRACH motif (D=[AUG], R=[AG], A, C, H=[AUC]), the main sequence recognition motif for the m6A writer enzyme, METTL3 (46) (**Figure 2D**). A site paired with an apparent m6A site overlaps with the m5C writer enzyme, NSUN6, sequence recognition motif, CUCCA (47). The second site overlaps with a sequence that closely resembles the NSUN6 motif, CUCCU. The third paired Nanocompore significant site didn’t closely resemble any known RNA modification writing sequence recognition motif. All six of these sites have detectable ionic current differences between the Native and IVT samples (**Figure 2E-2G**). In addition, they also separate into two distinct clusters in terms of ionic current and dwell time (Nanocompore GMM correct p-value < 0.01 and absolute value of the Log Odds ratio > 0.5, **Figure 2H-2J**).

This proximity of several Nanocompore significant sites suggests that there may be an association between significant sites on the same RNA molecules. To test this, we determined the modification status for each read at every Nanocompore significant site by using the probability of being assigned to the modified cluster in the GMM. Then we performed a chi-squared test of independence on the modified counts for each pair of Nanocompore significant sites on the same molecules and the expected modified counts under the assumption of independence (see methods). We found 28 sites that were significant for co-modification, 35 that were mutually exclusively modified, and 2 that were co-unmodified on the same molecules more than expected given the modification levels at each site (**Figure 3A and Supplementary Table 6**). The longest linear distance between co-modified sites was 392 nt found on RPS11-201, and the shortest distance was 5 nt, as on MT-ND1-201 (**Figure 3B**). Mitochondrial transcripts represented 21 of the 28 significant co-modified pairs of sites, and although mt-tRNA and mt-rRNA modifications have been characterized, far less is known about mt-mRNA modifications (48). Besides unclassified sites (42), m6A co-modified with m6A (10) are the most commonly co-modified inferred RNA modifications. This is unsurprising as m6A is the most abundant mammalian mRNA modification. However, m6A co-modified with m5C (4) is the second most common non-unclassified coordinated modification pair. This is unsurprising because three of the top ten enriched kmers across all the significant sites were DRACH motifs and four were associated with sequences enriched in miCLIP experiments for the m5C writer enzyme, NSUN6 (47) (**Figure 3C**). Although we only found evidence of four inferred m6A sites co-modified with inferred m5C sites on the same molecules, we found these modifications in proximity of each other in reference transcripts (**Figure 3D**). We found 7459 pairs of inferred m6A and inferred m5C sites on the same reference transcripts with an average distance of 287 nt between them, of which 1026 (13.76%) were within 25 nt each other. In contrast, we randomly shuffled ten times the Nanocompore significant sites within genes that we detected RNA modifications, and inferred the identity of these shuffled sites and determined the distance between the shuffled m6A and m5C sites. We found on average 5225 ± 103 pairs of sites that were on average 1587 ± 50 nt apart with 73 ± 11 (1.4% ± 0.22%) within 25 nt of each other. The proximity of the bulk inferred m6A and inferred m5C sites suggests that there is some shared function of these two RNA modifications. The lack of detection in single-molecule coordination tests is likely a result of our tests being under-powered to adequately detect coordination of RNA modifications on single-molecules with the number of reads we have for each transcript.

**Figure 3:**
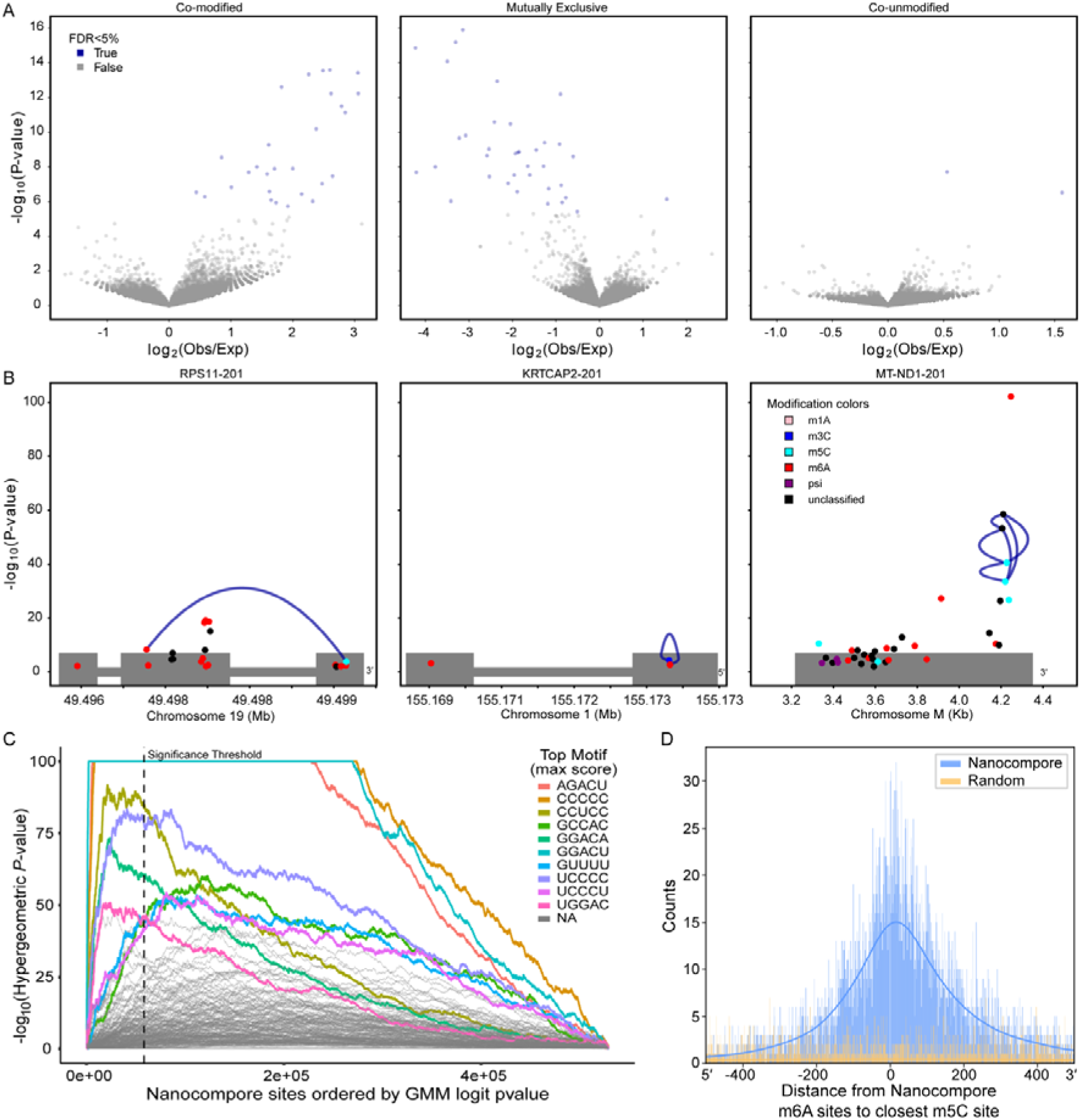
RNA modification co-occurance. (A) Volcano plots display pairwise co-modification patterns, mutual exclusivity, or co-unmodified status between single-molecule modified sites. Each dot corresponds to a pair of Nanocompore significant sites located on the same transcript. Blue dots represent statistically significant pairs at a 5% false discovery rate (FDR), while grey dots are non-significant. The vertical axis shows the –log10 of the Chi-squared test *p*-value, indicating statistical independence, and the horizontal axis represents the log2 ratio of observed versus expected co-occurrence counts. The transcript shown corresponds to the KATCAP2 isoform ENST00000295682 annotated in Gencode v36. (B) Three representative transcripts are shown. Each dot marks a site identified by Nanocompore as modified, with colors denoting the predicted modification type. The horizontal axis shows genomic position, and the Gencode v46 reference transcript structure is plotted in grey below, with strand direction (5′ or 3′ end) indicated. The vertical axis displays the Nanocompore GMM Logit –log10 *p*-value, and blue connecting lines highlight significant co-modified site pairs based on the Chi-squared test (FDR ≤ 5%). (C) A k-mer enrichment analysis ranks sequences associated with predicted Nanocompore modification sites. K-mers are ordered along the x-axis by their GMM-derived *p*-values, while the y-axis represents the –log10 of the hypergeometric *p*-value indicating enrichment. The ten highest-scoring k-mers are labeled, with sequences shown in the legend. If a k-mer is linked to known RNA modification types, this is noted in parentheses. Non-enriched k-mers are shown in grey (NA). (D) Histogram of genomic distances between predicted m6A and the nearest m5C Nanocompore sites within the same transcript isoform. The position “0” marks the m6A site, with negative values indicating upstream m5C sites and positive values downstream. Blue bars represent observed Nanocompore site distances, while yellow bars depict randomized controls.

### Differentially Modified Isoforms

We identified 1766 genes with at least one Nanocompore significant site. Among these, 1301 genes only had one isoform and 465 genes had multiple isoforms (**Figure 4A**). As expected, most isoforms only had a single significant site (**Figure 2A**). However, surprisingly, these sites often differed across isoforms of the same gene (**Figure 4B**). We found 9403 (35.3%) Nanocompore significant sites that were detected in a single isoform, despite sufficient coverage at the same genomic position in the other isoforms for Nanocompore to conduct the test (**Figure 4B**, purple region). However, 3810 (14.3%) of Nanocompore significant sites were shared with at least one other isoform at the same genomic position (**Figure 4B**, yellow region), and 11933 (44.8%) sites that were either found on an exon unique to that isoform (**Figure 4B** orange region) or 1472 (5.5%) sites that had insufficient coverage to test with Nanocompore (**Figure 4B** green region). Interestingly, in the case of PABPC1, a gene that binds poly(A) tails and is involved with translation and NMD (49), we detected a single position that was significant in 14 of the 19 different isoforms (**Figure 4A**). We were unable to identify the possible RNA modification at this site. The significant kmer sequence is UUUUG, and there are no annotated RNA modifications in proximity of the site.

**Figure 4:**
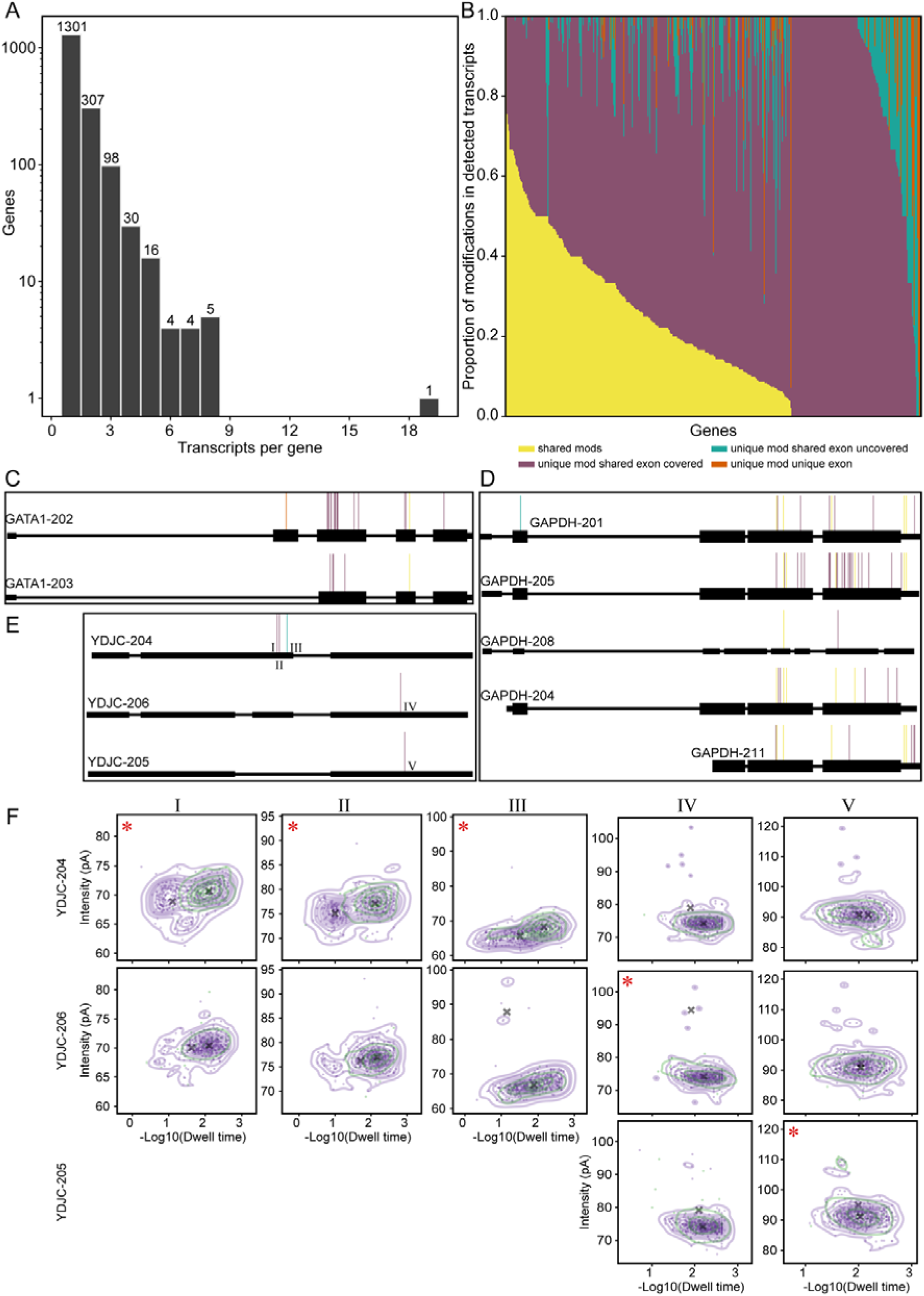
Isoform differential RNA modifications. (A) Histogram of the number of isoforms per gene detected with at least one Nanocompore significant site. (B) Genes with more than 1 reference transcript with a Nanocompore significant site. The regions are colored according to how the significant sites are shared among the detected transcripts. The x-axis is the genes, and the y-axis is the proportion of modifications shared among the different isoforms. Yellow represents modifications that were detected in at least 1 other isoform. Purple represents modifications detected in a single isoform, but there was sufficient coverage in at least 1 other isoform for Nanocompore to test for a modification. Green represents modifications detected in only 1 isoform, and there was insufficient coverage in any other isoform for Nanocompore to test for a modification. Orange represents modifications detected on an exonic region that does not exist in any other isoform. (C-E) Example gencode v36 transcript isoform models detected with at least one Nanocompore significant site for (C) GATA1, (D) GAPDH, and (E) YDJC. All transcript models are in the 5′-to-3′ orientation. The significant sites share the color coding as in (B). (E) has the five Nanocompore significant sites labeled I, II, III, IV, and V, which correspond to the panels in (F). (F) Ionic current distributions for the five Nanocompore significant sites found for YDJC. The y-axis is the ionic current intensity, and the x-axis is the –log10 of the dwell time. Reads from the Native sample (purple) and IVT sample (green) are shown. Black X’s indicate the GMM cluster centroids. The five sites are labeled in the upper left corner and a red astritic indicates that site was significant (Adjusted GMM logit pvalue <= 0.01 and Absolute value of the log odds ratio > 0.5).

The landscape of Nanocompore significant sites across different isoforms is complex, as can be seen in three representative genes, GATA1, GAPDH, and YDJC. In agreement with our observations across all isoforms, most of the Nanocompore significant sites are found in one of the isoforms (purple marks), with few shared among the isoforms (yellow marks) (**Figure 4C-4E**). A closer inspection of the ionic current data for YDJC isoforms shows that there is clear separation between the native and IVT reads for YDJC-204 at positions I, II, and III. YDJC-206, is not significant for a modification at these three positions and as expected, there is more overlap in the ionic current between the Native and IVT samples. Isoform YDJC-205 is lacking the exon for positions I, II, and III, and thus doesn’t have any ionic current alignments for those positions (**Figure 4F**). These results demonstrate the complexity of how RNA modifications are distributed throughout the transcriptome and allows for further investigations into isoform differential modifications and possible functional differences.

## Conclusions

In this study, we tested RNA in vitro transcription (IVT) to generate a modification-free transcriptome as a reference sample to detect RNA modifications using dRNA-seq. In our study, the IVT was limiting the analysis to a subset of the transcripts detected by the Native sample (**Figure 1C**). We can envision overcoming this limitation by either sequencing the IVT sample more deeply, or by altering the modification comparison framework to use the IVT data independent of the positions where the strands align but instead a pool of ionic current data akin to an ionic current kmer look-up table.

Despite the limitations of the IVT sample, we were still able to detect 26619 positions that were significant when compared with the Native sample (GMM logit FDR p-value <= 0.01 and the absolute value of the Log Odds Ratio > 0.5) (**Supplementary Table 4**). This is consistent with other attempts to identify RNA modifications using Nanocompore, which generally has higher specificity but lower sensitivity (50). Since the IVT sample is RNA modification-free, all RNA modifications are detected simultaneously, however, it is not possible to know the identity of the detected RNA modification from this analysis alone. As m6A is the most abundant internal mammalian mRNA modification (51), it was unsurprising that the Nanocompore significant sites we detected were distributed similarly to previously annotated m6A sites (43), with some differences (**Figure 2B**). We could infer that 41.13% significant sites were m6A and that 8.94% sites were m5C using known m6A and m5C writer sequence motifs and annotated m6A and m5C sites from RMBase3.0 (41) (**Supplementary Table 5**). It is important to note that our characterization of multiple modifications is indirect since our inferences rely on known sequence motifs, or previously annotated modification sites. Thus, when a site exhibits signal differences consistent with m6A or m5C, the final classification is contingent upon these external data. This limitation is shared by most comparative approaches and should be considered when interpreting results. We anticipate that continued improvements in these techniques, along with standardized computational frameworks, will enhance our ability to interrogate the epitranscriptome and elucidate the roles of individual RNA modifications in gene regulation.

Surprisingly, we found 7459 pairs of inferred m6A and m5C sites on the same reference transcripts, of which 1026 (13.76%) were within 25 nt each other **(Figure 3D**). We also found significant enrichment for four inferred m6A and m5C modifications on the same molecules for two transcripts, RPS11 and UBE25 (**Supplementary Table 6**). Recent advances in m5C detection methods are also finding that m5C is more prevalent in mRNA than previously thought (52,53). Furthermore, Mateos et. al. also found that m6A and m5C are in close proximity and co-occurring on the same molecules at higher than expected rates in poly(A) RNA samples from HEK293 and HeLa cells using a different nanopore RNA modification detection method (54).

It has been previously documented that m6A and m5C jointly promote methylation of each other, and together enhance protein translation of CDKN1A in response to stress agents (55). Although the mechanism for how m6A promotes m5C methylation and vice versa is not entirely clear, it is becoming apparent that the relationship between m6A and m5C is more widespread than previously thought as better m6A and m5C detection methods become available. Furthermore, there may be temporal components to the mechanism of methylation cooperation. There is a precedent for RNA modifications being deposited in successive order in rRNA biogenesis (56). However, it is unclear if the order of modification deposition is step-wise or stochastic at each stage of rRNA maturation (56). Simultaneous detection of different RNA modifications on single-molecules could help resolve potential temporal components of RNA modification deposition cooperation. We only detected evidence for four m6A sites coordinated with m5C sites on single molecules (**Supplemental table 6**), but 1026 such pairs of sites within 25 nt on the same reference transcripts (**Figure 3D**). This suggests that we were underpowered to detect the modifications on single molecules and with more coverage or better classification methods could better detect possible modification coordination on single-molecules if it exists.

Not only can long-reads from dRNA-seq be used to molecularly phase multiple RNA modifications, but also map efficiently to isoforms (57). This can be used to resolve isoform specific modification patterns (58). In this study, we found 1766 genes with Nanocompore significant sites, among which, 465 had significant sites in at least two different isoforms (**Figure 4**). The number of isoforms per gene that we found with Nanocompore significant sites ranged from two (307) to nineteen (1). Most of the sites were found in only one of the isoforms indicating that modification deposition was not consistent across isoforms. In some cases there is a hyper-modified isoform, and a hypo-modified isoform like seen in GATA1. In other cases, there was a mix of shared and unique significant sites, like seen in GAPDH (**Figure 4C-D**). And in a few cases, there were Nanocompore significant sites that were all unique to the different isoforms as seen in YDJC (**Figure 4E**).

There are several possible technical and biological reasons for detecting different patterns of RNA modifications in the different isoforms. One technical complication is the number of reads mapped to each isoform. The GATA1 isoform, GATA1-202, has more Nanocompore significant sites, but is also the dominant isoform detected with 10-fold more native reads than GATA1-203, 3613 and 342 native reads respectively (**Figure 4C**). For detection by Nanocompore, more than 2000 reads are suggested to detect a modification with 10% stoichiometry (35), and thus we likely miss several low stoichiometry modified sites due to lack of adequate coverage. Additionally, the IVT reads are not always evenly distributed between the various isoforms and thus certain sites in different isoforms may not be detectable due to a lack of IVT reads at that site.

One possible biological reason is the exon structure of the isoform. For example, the three YDJC isoforms we detected all have 470, 584, and 446 native reads respectively (**Figure 4E**). However, YDJC-204 is the only isoform significant for a modification at sites I, II, and III. Unsurprisingly, YDJC-205 has no reads that cover these sites because it lacks the exon.

YDJC-206 does have the exon, but was not significant at any of the three sites (59). Uzonyi et. al. recently proposed a model for m6A deposition, where the splicing machinery excludes the methylation machinery from accessing potential target sites for modification deposition (59). The only structural differences between YDJC-204 and YDJC-206 is an alternative TSS and an intron upstream of sites I, II, and III. The YDJC-206 upstream intron could be preventing the modification deposition. The ionic current distributions of the Native reads for YDJC-204 and YDJC-206 separate into two distinct clusters at the three sites whereas the IVT reads form a single cluster (**Figure 4F**). Despite the few possibly modified reads for YDJC-206, they are greatly reduced when compared to YDJC-204 and do not pass the significance threshold for our Nanocompore analysis. Based on visual inspection of the ionic current distributions, it is less clear why YDJC-206 is significant at site IV and YDJC-205 is significant at site V. It is possible that these two sites are false positives. Alternatively, the underlying RNA modification could have a more subtle effect on the ionic current that is detected by the GMM and not clear from visual inspection. Without a clear indication of which RNA modification is present at these two sites, it is difficult to resolve the truth at these two sites without a comprehensive RNA modification writer enzyme KD/KO screen.

All together, these results demonstrate the utility and challenges using an IVT transcriptome for detecting the positions of RNA modifications with dRNA-seq. Identifying different RNA modifications on the same molecules gives insights into possible coordination of different RNA modifications, and their distribution amongst different isoforms. Detecting these coordinated RNA modifications can be used to design follow-up experiments to determine the mechanism or function of these coordinated sites.

Furthermore, evaluating single-sample trained RNA modification detection models remains a challenging problem. As these methods become more widely adopted, repositories of IVT dRNA-seq datasets are invaluable true negative test cases to evaluate false positive rates. It is also unclear how quickly robust trained models can be produced for the nearly 170 known naturally occurring RNA modifications (22) and the nearly limitless possible synthetic RNA modifications (60). Until that time, comparing native dRNA-seq datasets to matched IVT dRNA-seq datasets permits detecting the multitude of known RNA modifications, and possibly highlighting possible locations to investigate further for potentially novel RNA modifications that wouldn’t be detected by these trained models.

In conclusion, IVT-based calling of RNA modifications currently represents the ideal strategy for deciphering the location of the multitude of modifications that make up the whole epitranscriptome, whose composition and dynamics remain to be investigated.

## Methods

### Cell culture Total RNA extraction and mRNA purification

K562 cells were grown in RPMI-1640 medium supplemented with 10% FBS (Sigma-. Aldrich F2442). Total RNA was extracted from 10 Million cells using Qiazol (Qiagen 79306) and RNeasy Micro Kit (Qiagen 74004). mRNA purification was performed with 100ug of Total RNA using µMACS™ mRNA Isolation Kit (Miltenyi Biotec 130-075-201) following the manufacturer’s protocol.

### cDNA synthesis protocol

Reactions for cDNA synthesis were prepared in 0.2 ml nuclease free PCR tubes on ice. First, 100 ng of mRNA were hybridized with 1 µM of the PolyT primer (5′-T_30_VN-3′) and 1 mM dNTPs in 6 µl. The samples were mixed gently by pipetting and centrifuged briefly. Then Incubated for 5 minutes at 70°C and left at 4°C until next use. Second, the hybridized RNA was mixed with 4 µl of RT buffer (1X Template Switching buffer, 3.75 µM Template Switching Oligo (5′-ACTCTAATACGACTCACTATAGGGAGAGGGCrGrGrG-3′), 1X Template switching RT Enzyme), and incubated at 42°C for 90 minutes, heat inactivated at 85°C for 5 minutes. The 10 µl of the first strand RT products were PCR amplified in 50 µl total volume of PCR buffer (1X Q5 Hot Start Master Mix (NEB #M0466S) and 0.2 µM T7 extension primer (5′-GCTCTAATACGACTCACTATAGG-3′)), with 15 cycles of 98°C for 10 sec, 68°C for 15 sec, and 72°C for 3 minutes. The amplified cDNA library was purified with AMPure beads (Beackman Coulter 10136224) following the manufacturer’s recommendations. Amplified cDNA was stored at –20°C for up to a week before use as the IVT template.

### RNA Synthesis

*In vitro* transcription was performed with the NEB HiScribe^®^ T7 High Yield RNA Synthesis Kit (NEB#E2040) in 20 µl of total volume, with 1ug of the previously prepared cDNA product mixed with final concentration 10 mM of: ATP, GTP, UTP, CTP, RNase Inhibitor (Thermofisher 10777019); and 2 µl of the T7 polymerase mix. The reaction was incubated at 42°C for 2 hours in a thermocycler.

### IVT RNA purification using phenol chloroform

The resulting IVT was purified by adding equal volume of phenol:chloroform:isoamyl alcohol (25:24:1) (Thermofisher 15593031) to the sample. It was Centrifuged for 5 minutes at 12,000 g at 4°C. The aqueous phase was extracted and transferred to a fresh tube. Equal volume of Chloroform was added to the fresh tube and was centrifuged for 5 minutes at 12,000 g at 4°C (this step was performed twice). The aqueous phase was transferred into a fresh tube with 2.5 volumes 100% ethanol and 0.1 volume 3M sodium acetate. Sample was incubated at –80°C for 1 hour, after that time it was centrifuged for 30 minutes at 12,000 g at 4°C to pellet the RNA. The supernatant was discarded and 500 ul of fresh 70% ethanol was added to the pellet. The sample was centrifuged for 5 minutes at 12,000 g at 4°C and the supernatant was discarded again (this step was repeated twice). The RNA was resuspended in 15ul of H20.

### Extended IVT RNA 3′ ends with poly(A)

A maximum of 10 ug of RNA were used for *in vitro* tailing. In 20 µl, the reagents were added in order 1-10 µg IVT RNA, Reaction Buffer to final concentration 1X, 1 mM ATP, 5 units of PolyA polymerase (*E. coli* or yeast as appropriate) (NEB#M0276S). The reaction was incubated at 37°C for either 7 or 30 minutes and stopped by purification with a MEGAclear column (Termofisher AM1908) following the manufacturer’s protocol. The purified RNA was optionally enriched for PolyA tails using a MultiMACS mRNA Isolation Kit (Miltenyi Biotec 130-075-201) following the manufacturer’s protocol.

### Direct RNA Nanopore Sequencing

RNA was adapted for sequencing using the standard SQK-RNA002 library protocol from ONT and sequenced on R9.4.1 MinION or PromethION flowcells. The resulting fast5 files were basecalled after sequencing with guppy v6.4.6 using the high accuracy model detailed in the configuration file for MinION (rna_r9.4.1_70bps_hac.cfg) or PromethION (rna_r9.4.1_70bps_hac_prom.cfg) flowcells, as appropriate.

### RNA modification detection

The fastq files for the IVT samples were concatenated into a single fastq file, and the directories containing the fast5 files were moved into a single directory to make indexing all the experiments together easier. This was also done for the native samples, but kept in a separate directory from the IVT samples. The two samples were processed as described previously (35). Briefly, the fastq files were aligned to the gencode reference transcriptome (v36) (61) concatenated with the SIRVs set 3 reference transcriptome (Lexogen 051.01) using minmap2 (v2.20) (62) and were filtered and sorted for primary forward alignments using samtools (v1.12) (63). The fast5 files were indexed and the ionic current events in the fast5 files were aligned to the transcriptome reference using nanopolish (0.13.3) (64). The resulting output was collapsed and the position-by-position pairwise modification detection was done using nanocompore (v1.0.4) (35) using context 0. To control for a single RNA modification being called in several neighboring sites, the Nanocompore –log10 transformed GMM pvalues for each transcript were peakcalled using scipy.signal.find_peaks with a floor of 2 (corresponding to a pvalue <= 0.01) and a distance of 5 (the kmer width). The resulting peakcalled sites were considered significant for a modification if the absolute value of the log odds ratio was also greater than 0.5 (35).

### Single Molecule Analysis

Normally, the cluster probabilities for each read is not linked to other positions in the transcript. To map the cluster probabilities at each position to the read of origin, the ionic current intensity and dwell times in the Nanocompore output db were matched to the ionic current intensities and dwell times in the eventalign_collapse.tsv file. In this way the read IDs could be linked to the GMM cluster probabilities. Then, the cluster with more Native sample label reads as the most probable cluster was selected as the “modified” cluster. A read was considered modified at any position if the modified cluster probability was greater than 0.75. The modification pattern was determined for all transcripts with two or more significant sites by creating a matrix with the number of significant site columns and number of read rows.

The matrix values were 0 when the site for that read was unmodified and 1 when modified. The observed counts were determined by counting all the instances of each unique pattern in the matrix. The expected counts were determined by calculating the linear combination of the observed frequencies at each site. The chi2 test of independence was computed using the observed and expected counts for each pattern of modification for each transcript assuming a uniform distribution. The resulting pvalues were multiple testing corrected using the Benjamini-Hochberg FDR procedure.

### Additional bioinformatics analysis

Unless otherwise noted, default conditions were used for all analysis tools. Transcriptomic coordinates were converted to genomic coordinates using bedparse (v0.2.3) (65). Genomic coordinates were translated to transcriptomic coordinates using R library GenomicFeatures function, mapToTranscripts (version 1.46.1). The EMBOSS function, fuzznuc (6.6.0.0) (66), was used to find the coordinates of possible METTL3, METTL16, and NSUN6 sequence recognition motifs in either genomic or transcriptomic coordinates. A Nanocompore significant site identity was inferred if the nanocompore 5-mer was within 2 nucleotides of a writer enzyme sequence recognition motif or an RMBase3.0 annotation (41) using bedtools closest (v2.29.1) (67). Bedtools closest was also used to determine distances between inferred m6A and m5C sites. Metagene plots were made using the R library, Guitar (2.10.0) (68). Nanocompore kmer enrichment analysis was done with Sylamar (v12-342) (69).

## Declarations

### Ethics approval and consent to participate

Not applicable.

### Consent for publication

Not applicable.

### Availability of data and materials

All data used in this study can be found at the European Nucleotide Archive using accession number PRJEB81473. Scripts used to analyze the data can be found at https://github.com/nicassiolab/K562_IVT.

## Author’s contributions

LM and LCT designed the study. LCT and CR performed the experiments, LM, SM, MF, TL, and PM performed the analysis. LCT and LM wrote the initial manuscript. EB, FN, MP, TL, MJM, TF supervised the project and edited the manuscript.

## Acknowledgements

We thank Diego Vozzi and the Genomics Facility at the Center for Human Technologies (CHT@Erzelli) of the Istituto Italiano di Tecnologia (IIT) and Clelia Peano the National Genomics Facility, Human Technopole for conducting the PromethION sequencing for this project. We thank Luca Rotta at the European Institute of Oncology (IEO) Genomic Unit for conducting the MinION sequencing for this project. We thank Daphne Welter and Vladimir Benes from EMBL genecore for sequencing support. We thank Salvatore Bianchi for managing all laboratory materials used in this study. We also thank Saul Pierotti for thoughtful discussions about appropriate statistical testing. Figure 1A was created with BioRender.com and released under a Creative Commons Attribution-NonCommercial-NoDerivs (CC-BY-NC-ND) 4.0 International license.

## Supplementary Results

### In vitro transcription optimization

We tested two Template Switching RT Enzymes (NEB and SmartScribe, see methods for details) using poly(A) RNA extracted from K562 cells (myelogenous leukemia cell line). The cDNA was assessed using qRT-PCR on three genes that cover a range of expression (MYC, RPPo, LNCRP) and a synthetic spikein (Lexongen SIRV MIX3) sequences (See methods).

The qRT-PCR for NEB ranged from 1 to 10 cycles lower than for SmartScribe **(Supplementary table 1**), and ERCC113 was not detected by SmartScribe where it was detected by NEB. The cDNA product from the NEB kit was used as the template for the IVT reaction. We found that the IVT incubation temperature had an effect on the quality of the IVT RNA (**Supplementary** Figure 2). The majority of transcripts were between 200 bp and 500 bp when incubated at 37°C, whereas the IVT strands ranged from 500 bp to 1000 bp when incubated at 42°C **(Supplementary** Figure 1). Both cDNA synthesis and IVT reduce the overall length of the samples when compared to total RNA and mRNA (**Supplementary** Figure 1). This is likely due to the processivity of the two enzymes used for these steps and something to consider carefully when producing IVT RNA from cellular RNA. IVT RNA quality is not the only aspect that can affect dRNA-seq throughput. The dRNA-seq library construction protocol requires a 3′ end poly(A) tail that is at least 10 nucleotides long. The full-length IVT constructs should include a 30 nucleotide long poly(A) tail, but incomplete synthesis products will be lacking this tail. However, the reference sample doesn’t need to perfectly match the structure of the sample transcriptome, so long as it has sufficient coverage across the transcriptome, regardless of fragment size. Thus, these incomplete synthesis products are still useful for ensuring coverage across each transcript.

To this end, we tested two different strategies to ensure proper library construction, *in vitro* polyadenylation (tailing), or combining tailing with poly(A) enrichment, and different methods to purify the IVT RNA prior to library construction (**Supplementary table 2**). We tested tailing the IVT RNA for 7 minutes with both the Yeast polyA polymerase (PAPY) and the E. coli polyA polymerase (PAPE) from NEB (see methods). We sequenced samples tailed from both enzymes in duplicate on MinIONs and collected 85,727 and 83,911 reads from the samples tailed with PAPY and 298,827 and 556,783 reads from the samples tailed with PAPE (**Supplementary table 2)**. Although the samples tailed with PAPE had higher throughput, all four samples had less throughput than expected. We increased the polyA tailing time from 7 minutes to 30 minutes with PAPE and included an additional phenol:chloroform:purification step after tailing in an attempt to further increase the throughput, as it has been shown that sequencing throughput is increased with cleaner RNA samples (28). These changes resulted in 947,857 reads from a MinION flowcell, indicating the protocol was optimized enough for PromethION flowcells.

## Supplementary Methods

### Cross platform modification detection

#### Data preprocessing

PromethION and MinION FAST5 files were basecalled with Guppy (v6) using the following commands respectively: *“guppy_basecaller –i fast5_directory –r –x ‘auto’ –s output_directory –-fast5_out –c rna_r9.4.1_70bps_hac_prom.cfg”, “guppy_basecaller –i fast5_directory –r –x ‘auto’ –s output_directory –-fast5_out –c rna_r9.4.1_70bps_hac.cfg”*. Reads were then aligned to the transcriptome with minimap2 (v2.17), and the resulting BAM files were processed with samtools (v1.6) to filter unmapped reads, not primary alignments, supplementary alignments, and alignments on the reverse strand: *“minimap2 –ax map-ont –k 14 reference_transcriptome.fasta reads.fastq | samtools view –h –F2324 | samtools sort –o filtered_reads_mapping_on_transcriptome_F2324.bam”*.

Filtered BAM files were imported in R with the *pileup* function of Rsamtools package (v2.14.0) to compute the number of nucleotides on transcripts 3′ UTRs. The minimum of this quantity across conditions for each transcript was used to select the 50 transcripts with higher 3′UTR signal for further analyses (see the R script *selection_transcripts.R*). Specifically, we applied Nanocompore (v1.0.4) to this dataset to call RNA modifications on K562 untreated samples, sequenced either on MinION or PromethION flow-cells, using IVT data acquired on the PromethION machine as baseline.

### RNA modifications profiling

Reads mapping to the 3′ UTR of the 50 selected transcripts were extracted using samtools (v1.6) and seqtk (v1.3) with the following commands: *“samtools view filtered_reads_mapping_on_transcriptome_F2324.bam –L bed_3utr_top50_tx.bed | cut –f1 | sort| uniq > reads_3UTR_50tx.txt”* followed by *“seqtk subseq reads.fastq reads_3UTR_50tx.txt > reads_3UTR_50tx.fastq”*. Then, the Nanocompore pipeline was run on these sub-sampled FASTQ files using minimap2 (v2.17), samtools (v1.6), f5c (v0.7) and Nanocompore (v1.0.4). See the scripts *nanocompore_script1.sh* and *nanocompore_script2.sh*.

### Nanocompore post-processing

Nanocompore output files (outnanocompore_results.tsv) obtained comparing the WT sample – sequenced either on MinION or on PromethION flow-cells – to the PromethION IVT baseline were post-processed in R: see the *post_processing_analysis_IVT.R* script.

### Fixed coverage

The aforementioned analysis was repeated on a random set of reads obtained capping the coverage on the 3′ UTR of the 50 selected transcripts to 100x with the samNormalise.pl script (see https://gitlab.com/gringer/bioinfscripts/-/blob/master/samNormalise.pl) and the following commands: *“minimap2 –ax map-ont –k 14 reference_transcriptome.fasta reads_3UTR_50tx.fastq | samtools view –h –F2324 | samtools sort –o filtered_reads_3UTR_50tx_mapping_on_transcriptome_F2324.bam”; “samtools view filtered_reads_3UTR_50tx_mapping_on_transcriptome_F2324.bam | samNormalise.pl – coverage 100 –format fastq > reads_3UTR_50tx_cov100.fastq”*.

## Supplemental Figures

**Supplementary Figure 1.**
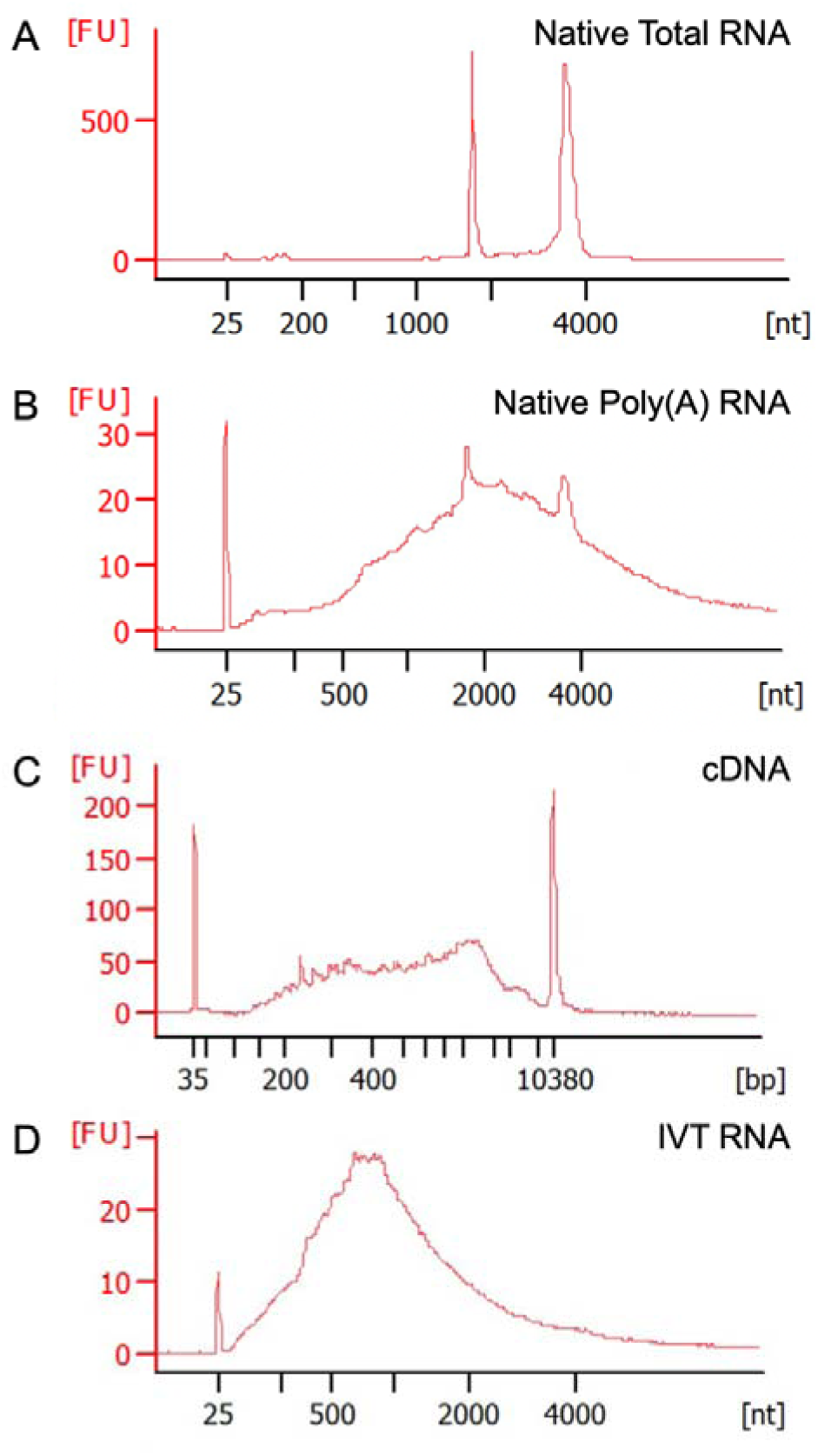
Electropherograms obtained with Bioanalyzer 2100 in K562 cells. Fluorescence intensity is represented in the y-axis and RNA size in nucleotides in the x-axis. (A) Profile of Total RNA loaded in the Nano-Chip. The two distinct sharp peaks of RNA represent the 18s (around 2000 bp ladder marker) and 28s (around 4000 bp ladder marker) and show evidence of high quality RNA. (B) Profile of mRNA loaded after polyA purification in the Pico-Chip. The profile displays reduction of ribosomal peaks corresponding to the mRNA distribution. (C) Profile of cDNA generated from k562 mRNA loaded on HighSensitivity-Chip. (D) Profile of IVT RNA generated from K562 cDNA after in vitro Polyadenylation loaded on Nano-Chip showing an average size of 1000bp.

**Supplementary Figure 2:**
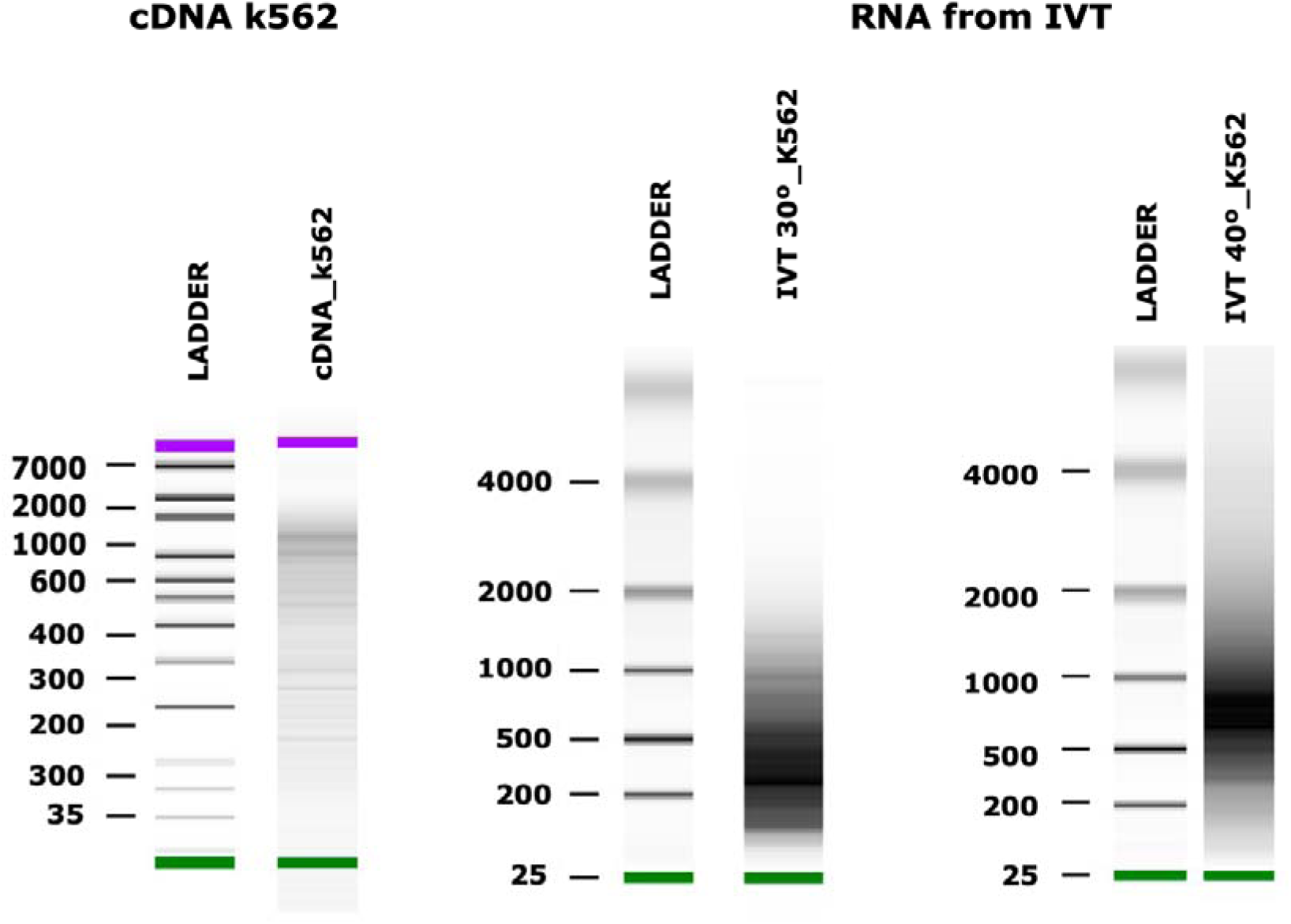
Bioanalyzer gel showing on the left the cDNA synthetized with template switching RT enzyme. On the right the RNA in vitro transcribed generated with 30° or 40° incubation temperatures.

**Supplementary Figure 3:**
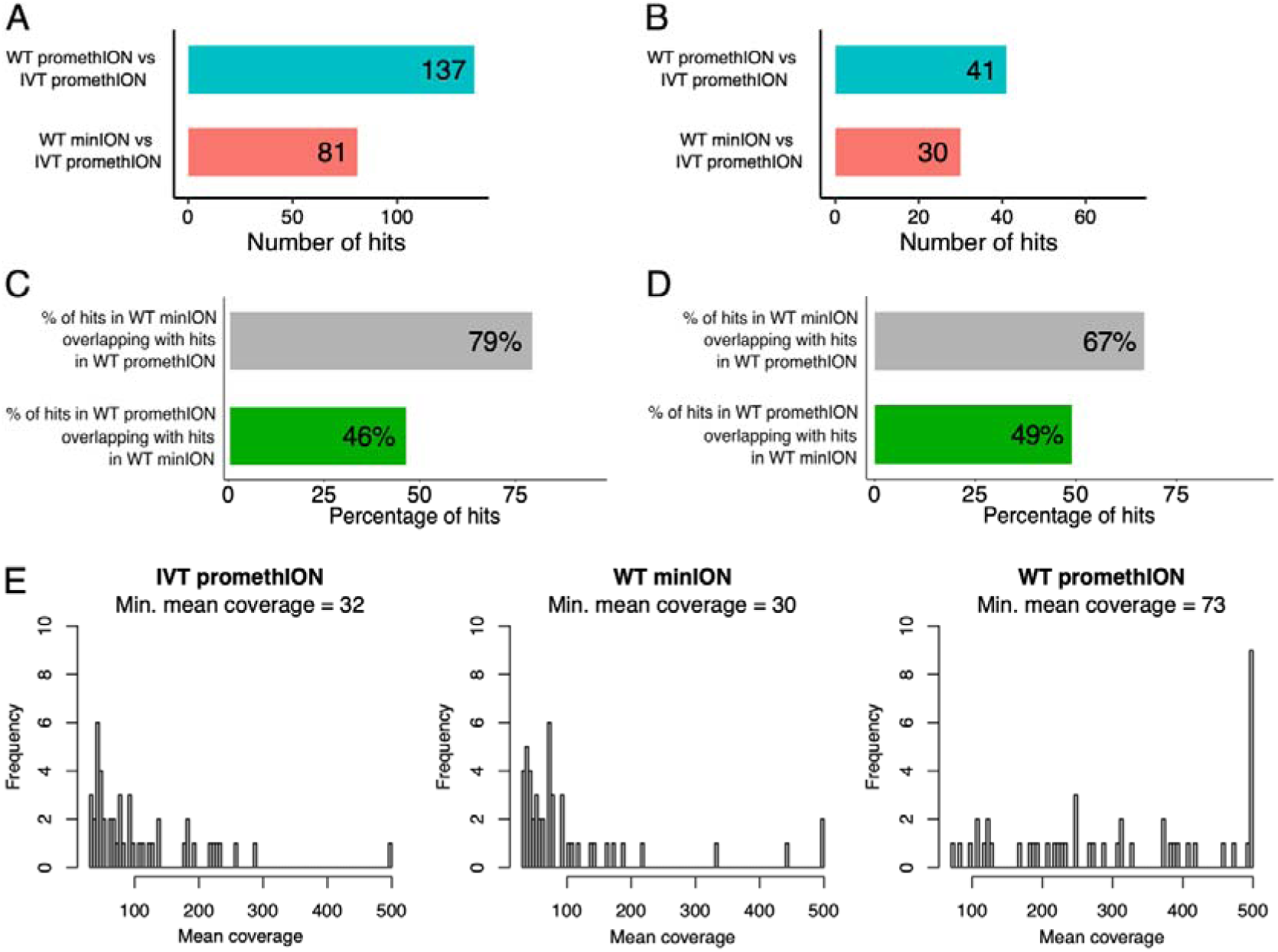
Comparison of MinION and PromethION Sequencing for RNA Modifications Profiling. (A) Number of hits identified by Nanocompore in the WT MinION (red) and PromethION (light blue) samples using the same IVT PromethION sample as a reference. (B) Same as (A) but capping the 3’ UTRs coverage to 100x. (C) Overlap between the hits identified by Nanocompore in the WT MinION sample and those detected in the WT PromethION sample using the same IVT PromethION sample as a reference. The size of the intersection is normalized to the number of hits detected in the MinION (gray) and PromethION (green) analyses, respectively. (D) Same as (C) but capping the 3’ UTRs coverage to 100x. (G) Distribution of the mean coverage considering only the sites analyzable in all the samples (saturation at 500x). All analyses were performed on 50 transcripts selected according to their 3’ UTRs coverage.

**Supplementary Figure 4:**
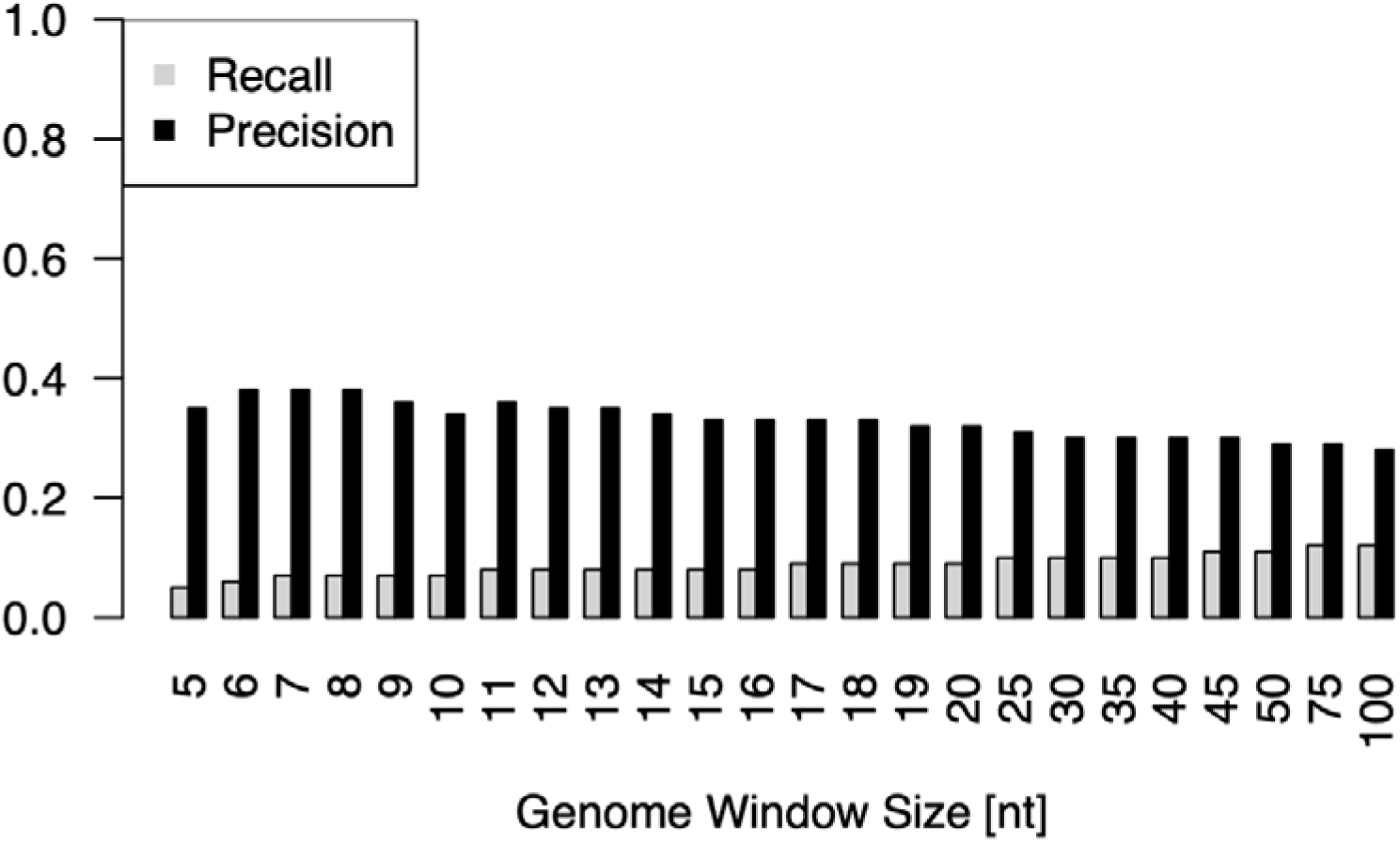
Precision and recall of Nanocompore-inferred m6A sites across different genomic bin sizes. A bin is considered detected if it contains at least one significant Nanocompore site, and considered modified if it contains at least one GLORI-seq hit. The analysis includes only bins derived from genes analyzed by Nanocompore and containing at least one RRACH motif.

**Supplementary Figure 5:**
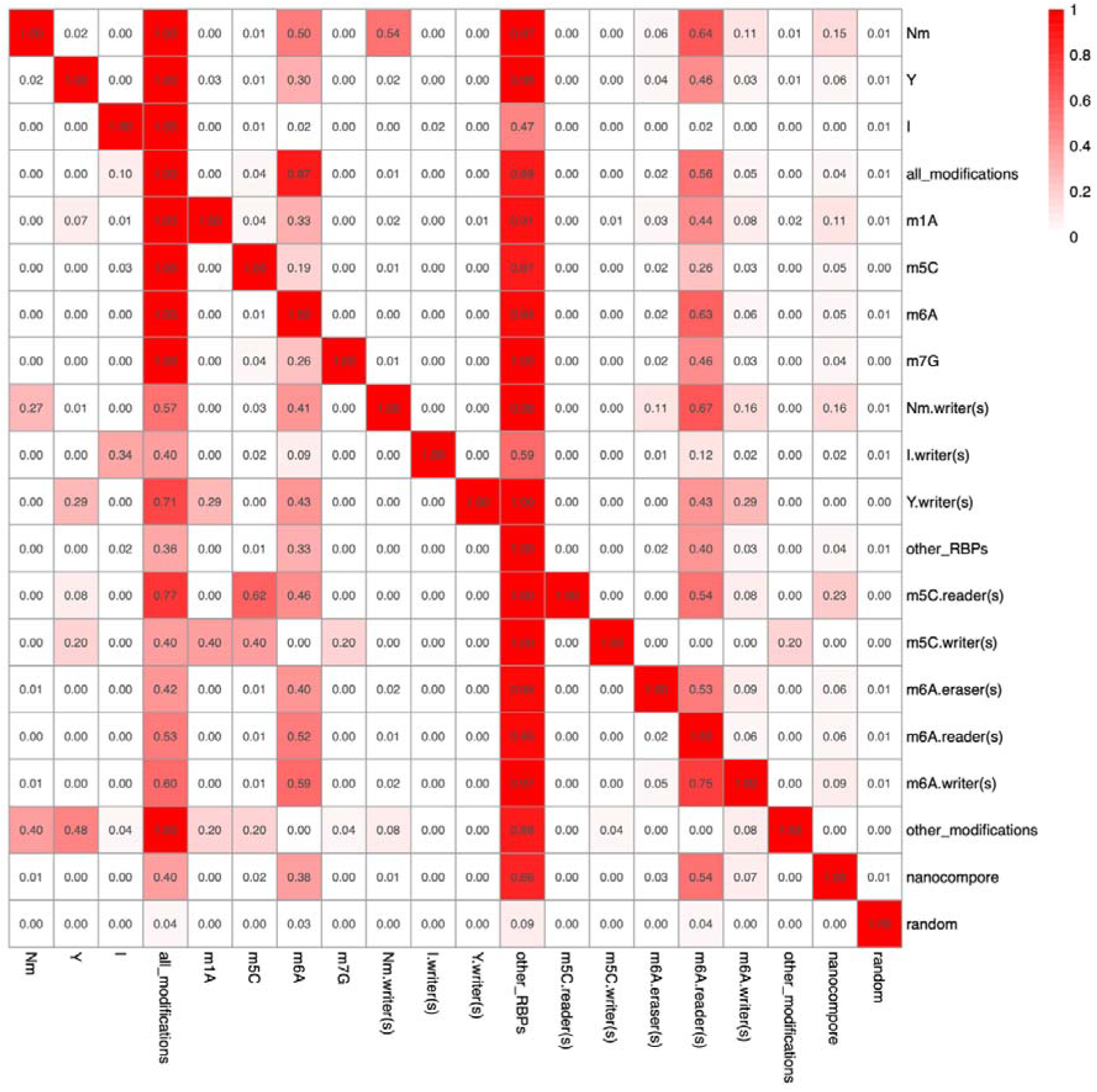
Full Heatmap reporting the overlap of the Nanocompore significant sites with every feature annotated in RMBase3.0. Figure 2C is a subset of this heatmap. The value in each cell of the matrix is calculated by dividing the intersection between the positive row and column bins divided by the total number of positive row bins. See diagram for reading the values in each cell.

## Supplementary Table legends

**Supplementary Table 1:** qRT PCR cycles of different genes and Spikein sequences using Template Switching RT or SmartScribe for cDNA synthesis.

**Supplementary Table 2:** Table describing the different Nanopore runs using GridiON and the different methods of RNA tailing and purification.

**Supplementary Table 3:** PromethION RNA002 sequencing summary statistics.

**Supplementary Table 4:** Nanocompore RNA modification detection summary statistics.

**Supplementary Table 5:** Modification inference summary using RMBase and writer motifs.

**Supplementary Table 6:** Significant single-molecule co-modification summary statistics and annotations.

## References

1. Acera Mateos P, J Sethi A, Ravindran A, Srivastava A, Woodward K, Mahmud S, Kanchi M, Guarnacci M, Xu J, W S Yuen Z, et al. 2024. Prediction of m6A and m5C at single-molecule resolution reveals a transcriptome-wide co-occurrence of RNA modifications. Nat Commun 15: 3899.

2. Anreiter I, Mir Q, Simpson JT, Janga SC, Soller M. 2021. New Twists in Detecting mRNA Modification Dynamics. Trends Biotechnol 39: 72–89.

3. Baylin SB, Jones PA. 2011. A decade of exploring the cancer epigenome — biological and translational implications. Nat Rev Cancer 11: 726–734.

1. Baylin SB, Jones PA. A decade of exploring the cancer epigenome — biological and translational implications. Nat Rev Cancer. 2011 Oct;11(10):726–34.

2. Esteller M. Non-coding RNAs in human disease. Nat Rev Genet. 2011 Dec;12(12):861–74.

3. Henninger JE, Oksuz O, Shrinivas K, Sagi I, LeRoy G, Zheng MM, et al. RNA-Mediated Feedback Control of Transcriptional Condensates. Cell. 2021 Jan;184(1):207–225.e24.

4. Fu XD, Ares M. Context-dependent control of alternative splicing by RNA-binding proteins. Nat Rev Genet. 2014 Oct;15(10):689–701.

5. Gebauer F, Hentze MW. Molecular mechanisms of translational control. Nat Rev Mol Cell Biol. 2004 Oct;5(10):827–35.

6. Wan Y, Qu K, Zhang QC, Flynn RA, Manor O, Ouyang Z, et al. Landscape and variation of RNA secondary structure across the human transcriptome. Nature. 2014 Jan 30;505(7485):706–9.

7. Roundtree IA, Evans ME, Pan T, He C. Dynamic RNA Modifications in Gene Expression Regulation. Cell. 2017 Jun;169(7):1187–200.

8. Gilbert WV, Bell TA, Schaening C. Messenger RNA modifications: Form, distribution, and function. Science. 2016 Jun 17;352(6292):1408–12.

9. Delaunay S, Helm M, Frye M. RNA modifications in physiology and disease: towards clinical applications. Nat Rev Genet. 2024 Feb;25(2):104–22.

10. Harcourt EM, Kietrys AM, Kool ET. Chemical and structural effects of base modifications in messenger RNA. Nature. 2017 Jan 19;541(7637):339–46.

11. Wang X, Lu Z, Gomez A, Hon GC, Yue Y, Han D, et al. N6-methyladenosine-dependent regulation of messenger RNA stability. Nature. 2014 Jan;505(7481):117–20.

12. Xiao W, Adhikari S, Dahal U, Chen YS, Hao YJ, Sun BF, et al. Nuclear m 6 A Reader YTHDC1 Regulates mRNA Splicing. Mol Cell. 2016 Feb;61(4):507–19.

13. Wang X, Zhao BS, Roundtree IA, Lu Z, Han D, Ma H, et al. N6-methyladenosine Modulates Messenger RNA Translation Efficiency. Cell. 2015 Jun;161(6):1388–99.

14. Schumann U, Zhang HN, Sibbritt T, Pan A, Horvath A, Gross S, et al. Multiple links between 5-methylcytosine content of mRNA and translation. BMC Biol. 2020 Dec;18(1):40.

15. Motorin Y, Helm M. tRNA Stabilization by Modified Nucleotides. Biochemistry. 2010 Jun 22;49(24):4934–44.

16. Kierzek E, Malgowska M, Lisowiec J, Turner DH, Gdaniec Z, Kierzek R. The contribution of pseudouridine to stabilities and structure of RNAs. Nucleic Acids Res. 2014 Mar 1;42(5):3492–501.

17. Monaco PL, Marcel V, Diaz JJ, Catez F. 2′-O-Methylation of Ribosomal RNA: Towards an Epitranscriptomic Control of Translation? Biomolecules. 2018 Oct 3;8(4):106.

18. Jady BE. A small nucleolar guide RNA functions both in 2’-O-ribose methylation and pseudouridylation of the U5 spliceosomal RNA. EMBO J. 2001 Feb 1;20(3):541–51.

19. Morais P, Adachi H, Yu YT. The Critical Contribution of Pseudouridine to mRNA COVID-19 Vaccines. Front Cell Dev Biol. 2021 Nov 4;9:789427.

20. Svitkin YV, Gingras AC, Sonenberg N. Membrane-dependent relief of translation elongation arrest on pseudouridine– and N1-methyl-pseudouridine-modified mRNAs. Nucleic Acids Res. 2022 Jul 22;50(13):7202–15.

21. Pardi N, Krammer F. mRNA vaccines for infectious diseases — advances, challenges and opportunities. Nat Rev Drug Discov [Internet]. 2024 Oct 4 [cited 2024 Oct 7]; Available from: https://www.nature.com/articles/s41573-024-01042-y

22. Boccaletto P, Machnicka MA, Purta E, Piątkowski P, Bagiński B, Wirecki TK, et al. MODOMICS: a database of RNA modification pathways. 2017 update. Nucleic Acids Res. 2018 Jan 4;46(D1):D303–7.

23. National Academies of Sciences, Engineering, and Medicine; Health and Medicine Division; Division on Earth and Life Studies; Board on Health Sciences Policy; Board on Life Sciences; Toward Sequencing and Mapping of RNA Modifications Committee. Charting a Future for Sequencing RNA and Its Modifications: A New Era for Biology and Medicine [Internet]. Washington (DC): National Academies Press (US); 2024 [cited 2024 Oct 7]. Available from: http://www.ncbi.nlm.nih.gov/books/NBK606036/

24. Kirpekar F, Douthwaite S, Roepstorff P. Mapping posttranscriptional modifications in 5S ribosomal RNA by MALDI mass spectrometry. RNA. 2000 Feb;6(2):296–306.

25. Linder B, Grozhik AV, Olarerin-George AO, Meydan C, Mason CE, Jaffrey SR. Single-nucleotide-resolution mapping of m6A and m6Am throughout the transcriptome. Nat Methods. 2015 Aug;12(8):767–72.

26. Carlile TM, Rojas-Duran MF, Zinshteyn B, Shin H, Bartoli KM, Gilbert WV. Pseudouridine profiling reveals regulated mRNA pseudouridylation in yeast and human cells. Nature. 2014 Nov;515(7525):143–6.

27. Garalde DR, Snell EA, Jachimowicz D, Sipos B, Lloyd JH, Bruce M, et al. Highly parallel direct RNA sequencing on an array of nanopores. Nat Methods. 2018 Mar;15(3):201–6.

28. Workman RE, Tang AD, Tang PS, Jain M, Tyson JR, Razaghi R, et al. Nanopore native RNA sequencing of a human poly(A) transcriptome. Nat Methods. 2019 Dec;16(12):1297–305.

29. Furlan M, Delgado-Tejedor A, Mulroney L, Pelizzola M, Novoa EM, Leonardi T. Computational methods for RNA modification detection from nanopore direct RNA sequencing data. RNA Biol. 2021 Oct 15;18(sup1):31–40.

30. Kovaka S, Hook PW, Jenike KM, Shivakumar V, Morina LB, Razaghi R, et al. Uncalled4 improves nanopore DNA and RNA modification detection via fast and accurate signal alignment. Nat Methods. 2025 Apr;22(4):681–91.

31. Delgado-Tejedor A, Medina R, Begik O, Cozzuto L, López J, Blanco S, et al. Native RNA nanopore sequencing reveals antibiotic-induced loss of rRNA modifications in the A– and P-sites. Nat Commun. 2024 Nov 29;15(1):10054.

32. Manrao EA, Derrington IM, Laszlo AH, Langford KW, Hopper MK, Gillgren N, et al. Reading DNA at single-nucleotide resolution with a mutant MspA nanopore and phi29 DNA polymerase. Nat Biotechnol. 2012 Apr;30(4):349–53.

33. Maestri S, Furlan M, Mulroney L, Coscujuela Tarrero L, Ugolini C, Dalla Pozza F, et al. Benchmarking of computational methods for m6A profiling with Nanopore direct RNA sequencing. Brief Bioinform. 2024 Mar 1;25(2):bbae001.

34. Begik O, Lucas MC, Pryszcz LP, Ramirez JM, Medina R, Milenkovic I, et al. Quantitative profiling of pseudouridylation dynamics in native RNAs with nanopore sequencing. Nat Biotechnol. 2021 Oct;39(10):1278–91.

35. Leger A, Amaral PP, Pandolfini L, Capitanchik C, Capraro F, Miano V, et al. RNA modifications detection by comparative Nanopore direct RNA sequencing. Nat Commun. 2021 Dec 10;12(1):7198.

36. Tavakoli S, Nabizadeh M, Makhamreh A, Gamper H, McCormick CA, Rezapour NK, et al. Semi-quantitative detection of pseudouridine modifications and type I/II hypermodifications in human mRNAs using direct long-read sequencing. Nat Commun. 2023 Jan 19;14(1):334.

37. Zhang Z, Chen T, Chen HX, Xie YY, Chen LQ, Zhao YL, et al. Systematic calibration of epitranscriptomic maps using a synthetic modification-free RNA library. Nat Methods. 2021 Oct;18(10):1213–22.

38. Song LF, Deng ZH, Gong ZY, Li LL, Li BZ. Large-Scale de novo Oligonucleotide Synthesis for Whole-Genome Synthesis and Data Storage: Challenges and Opportunities. Front Bioeng Biotechnol. 2021 Jun 22;9:689797.

39. McCormick CA, Akeson S, Tavakoli S, Bloch D, Klink IN, Jain M, et al. Multicellular, IVT-derived, unmodified human transcriptome for nanopore-direct RNA analysis. GigaByte Hong Kong China. 2024;2024:gigabyte129.

40. Wongsurawat T, Jenjaroenpun P, Anekwiang P, Arigul T, Thongrattana W, Jamshidi Parsian A, et al. Exploiting nanopore sequencing for characterization and grading of *IDH* mutant gliomas. Brain Pathol. 2024 Jan;34(1):e13203.

41. Xuan J, Chen L, Chen Z, Pang J, Huang J, Lin J, et al. RMBase v3.0: decode the landscape, mechanisms and functions of RNA modifications. Nucleic Acids Res. 2024 Jan 5;52(D1):D273–84.

42. Batista PJ. The RNA Modification N6-methyladenosine and Its Implications in Human Disease. Genomics Proteomics Bioinformatics. 2017 Jun;15(3):154–63.

43. Meyer KD, Saletore Y, Zumbo P, Elemento O, Mason CE, Jaffrey SR. Comprehensive analysis of mRNA methylation reveals enrichment in 3’ UTRs and near stop codons. Cell. 2012 Jun 22;149(7):1635–46.

44. Liu C, Sun H, Yi Y, Shen W, Li K, Xiao Y, et al. Absolute quantification of single-base m6A methylation in the mammalian transcriptome using GLORI. Nat Biotechnol. 2023 Mar;41(3):355–66.

45. Chandrasekhar C, Kumar PS, Sarma PVGK. Novel mutations in the kinase domain of BCR-ABL gene causing imatinib resistance in chronic myeloid leukemia patients. Sci Rep. 2019 Feb 20;9(1):2412.

46. Dominissini D, Moshitch-Moshkovitz S, Schwartz S, Salmon-Divon M, Ungar L, Osenberg S, et al. Topology of the human and mouse m6A RNA methylomes revealed by m6A-seq. Nature. 2012 May 10;485(7397):201–6.

47. Selmi T, Hussain S, Dietmann S, Heiß M, Borland K, Flad S, et al. Sequence– and structure-specific cytosine-5 mRNA methylation by NSUN6. Nucleic Acids Res. 2021 Jan 25;49(2):1006–22.

48. Boughanem H, Böttcher Y, Tomé-Carneiro J, López de Las Hazas MC, Dávalos A, Cayir A, et al. The emergent role of mitochondrial RNA modifications in metabolic alterations. Wiley Interdiscip Rev RNA. 2023 Mar;14(2):e1753.

49. Peixeiro I, Inácio Â, Barbosa C, Silva AL, Liebhaber SA, Romão L. Interaction of PABPC1 with the translation initiation complex is critical to the NMD resistance of AUG-proximal nonsense mutations. Nucleic Acids Res. 2012 Feb;40(3):1160–73.

50. Zhong ZD, Xie YY, Chen HX, Lan YL, Liu XH, Ji JY, et al. Systematic comparison of tools used for m6A mapping from nanopore direct RNA sequencing. Nat Commun. 2023 Apr 5;14(1):1906.

51. Anreiter I, Mir Q, Simpson JT, Janga SC, Soller M. New Twists in Detecting mRNA Modification Dynamics. Trends Biotechnol. 2021 Jan;39(1):72–89.

52. Hartstock K, Kueck NA, Spacek P, Ovcharenko A, Hüwel S, Cornelissen NV, et al. MePMe-seq: antibody-free simultaneous m6A and m5C mapping in mRNA by metabolic propargyl labeling and sequencing. Nat Commun. 2023 Nov 7;14(1):7154.

53. Lu L, Zhang X, Zhou Y, Shi Z, Xie X, Zhang X, et al. Base-resolution m5C profiling across the mammalian transcriptome by bisulfite-free enzyme-assisted chemical labeling approach. Mol Cell. 2024 Aug;84(15):2984–3000.e8.

54. Acera Mateos P, J Sethi A, Ravindran A, Srivastava A, Woodward K, Mahmud S, et al. Prediction of m6A and m5C at single-molecule resolution reveals a transcriptome-wide co-occurrence of RNA modifications. Nat Commun. 2024 May 9;15(1):3899.

55. Li Q, Li X, Tang H, Jiang B, Dou Y, Gorospe M, et al. NSUN2-Mediated m5C Methylation and METTL3/METTL14-Mediated m6A Methylation Cooperatively Enhance p21 Translation. J Cell Biochem. 2017 Sep;118(9):2587–98.

56. Sloan KE, Warda AS, Sharma S, Entian KD, Lafontaine DLJ, Bohnsack MT. Tuning the ribosome: The influence of rRNA modification on eukaryotic ribosome biogenesis and function. RNA Biol. 2017 Sep 2;14(9):1138–52.

57. Pardo-Palacios FJ, Wang D, Reese F, Diekhans M, Carbonell-Sala S, Williams B, et al. Systematic assessment of long-read RNA-seq methods for transcript identification and quantification. Nat Methods. 2024 Jul;21(7):1349–63.

58. Gleeson J, Madugalle SU, McLean C, Bredy TW, De Paoli-Iseppi R, Clark MB. Isoform-level profiling of m6A epitranscriptomic signatures in human brain [Internet]. 2024 [cited 2024 Oct 4]. Available from: http://biorxiv.org/lookup/doi/10.1101/2024.01.31.578088

59. Uzonyi A, Dierks D, Nir R, Kwon OS, Toth U, Barbosa I, et al. Exclusion of m6A from splice-site proximal regions by the exon junction complex dictates m6A topologies and mRNA stability. Mol Cell. 2023 Jan 19;83(2):237–251.e7.

60. Stephenson W, Razaghi R, Busan S, Weeks KM, Timp W, Smibert P. Direct detection of RNA modifications and structure using single-molecule nanopore sequencing. Cell Genomics. 2022 Feb;2(2):100097.

61. Frankish A, Carbonell-Sala S, Diekhans M, Jungreis I, Loveland JE, Mudge JM, et al. GENCODE: reference annotation for the human and mouse genomes in 2023. Nucleic Acids Res. 2023 Jan 6;51(D1):D942–9.

62. Li H. Minimap2: pairwise alignment for nucleotide sequences. Birol I, editor. Bioinformatics. 2018 Sep 15;34(18):3094–100.

63. Li H, Handsaker B, Wysoker A, Fennell T, Ruan J, Homer N, et al. The Sequence Alignment/Map format and SAMtools. Bioinforma Oxf Engl. 2009 Aug 15;25(16):2078– 9.

64. Simpson JT, Workman RE, Zuzarte PC, David M, Dursi LJ, Timp W. Detecting DNA cytosine methylation using nanopore sequencing. Nat Methods. 2017 Apr;14(4):407–10.

65. Leonardi T. Bedparse: feature extraction from BED files. J Open Source Softw. 2019 Feb 28;4(34):1228.

66. Rice P, Longden I, Bleasby A. EMBOSS: The European Molecular Biology Open Software Suite. Trends Genet. 2000 Jun;16(6):276–7.

67. Quinlan AR, Hall IM. BEDTools: a flexible suite of utilities for comparing genomic features. Bioinformatics. 2010 Mar 15;26(6):841–2.

68. Cui X, Wei Z, Zhang L, Liu H, Sun L, Zhang SW, et al. Guitar: An R/Bioconductor Package for Gene Annotation Guided Transcriptomic Analysis of RNA-Related Genomic Features. BioMed Res Int. 2016;2016:1–8.

69. Van Dongen S, Abreu-Goodger C, Enright AJ. Detecting microRNA binding and siRNA off-target effects from expression data. Nat Methods. 2008 Dec;5(12):1023–5.

